# Image-based patch selection for deep learning to improve automated Gleason grading in histopathological slides

**DOI:** 10.1101/2020.09.26.314989

**Authors:** William Speier, Jiayun Li, Wenyuan Li, Karthik Sarma, Corey Arnold

## Abstract

Automated Gleason grading can be a valuable tool for physicians when assessing risk and planning treatment for prostate cancer patients. Semantic segmentation provides pixel-wise Gleason predictions across an entire slide, which can be more informative than classification of pre-selected homogeneous regions. Deep learning methods can automatically learn visual semantics to accomplish this task, but training models on whole slides is impractical due to large image sizes and scarcity of fully annotated data. Patch-based methods can alleviate these problems, and have been shown to produce significant results in histopathology segmentation. However, the irregular contours of biopsies on slides makes performance highly dependent on patch selection. In the traditional grid-based strategy, many patches lie on biopsy boundaries, reducing segmentation accuracy due to a reduction in contextual information. In this paper, we propose an automatic patch selection process based on image features. This algorithm segments the biopsy and aligns patches based on the tissue contour to maximize the amount of contextual information in each patch. This method was used to generate patches for a fully convolutional network to segment high grade, low grade, and benign tissue from a set of 59 histopathological slides, and results were compared against manual physician labels. We show that using our image-based patch selection algorithm results in a significant improvement in segmentation accuracy over the traditional grid-based approach. Our results suggest that informed patch selection can be a valuable addition to an automated histopathological analysis pipeline.

## 1 Introduction

In the United States, prostate cancer is the third deadliest and most common cancer in men [1]. Prostate biopsy is one of the key components in monitoring patients with low to intermediate risk for clinically localized prostate cancer [2, 3]. Biopsies are performed repeatedly, each producing several tissue slides. Pathologists then scan through slides, searching for relevant regions with which to assign Gleason grades. This process can be time-consuming and tedious. Moreover, a single digital whole slide image (WSI) contains over a million pixels, which makes it difficult to analyze the entire slide at the highest resolution. Computational tools, which detect regions of interest (ROIs) on large-scale WSIs, can not only be used as a pre-step for fine-grained analysis such as biomarker discovery, but can also aid pathologists to quickly locate important tissue areas and potentially reduce the diagnosis time. Despite the potential impact, it is not trivial to develop an automatic cancer detection model due to the large variance of tissue appearances even within a single cancer grade as well as image artifacts caused by markers, dust, staining, etc.

Machine learning models are widely used for classification problems in the healthcare domain because of their ability to learn and utilize patterns from data to make predications [4]. Recent, developments in deep learning models [5] have drawn significant research interest because of their ability to automatically learn features specific to the data for classification and prediction tasks, achieving state-of-the-art performance in challenging problems. In particular, convolutional neural networks have been used for image classification and object detection (e.g., ImageNet [6], video classification [7], etc.).

A difficulty in medical imaging is that image sizes are often too large to train at once, and data scarcity limits the number of features that can be learned. A common approach to handling these issues is to use image patches, which has been successful in many anatomical domains, including brain [8] and prostate [9]. In biopsy slides, however, the majority of the image consists of whitespace or annotation, so patch selection is crucial for accurate training and segmentation. Traditional grid-based selection methods are suboptimal because the irregular shape of biopsy specimens results in large amounts of white space in these patches, which limits the amount of context available for a classification and reduces performance. The goal of this paper is to propose a method that uses the biopsy contour to select optimal patches for deep learning models. We evaluate this method by using it to provide for a previously published Path-CNN framework [10]. Results are compared against those achieved using a traditional grid-based patch selection.

## 2 Methods

The methods are described as follows: 1) first, the proposed patch selection method is described; 2) then, we outline the architecture for the deep learning classifier used for this segmentation task; and 3) the dataset used for evaluating the performance of these methods is described as well as the evaluation metrics and statistical tests that are used to evaluate performance.

### 2.1 Patch Selection

The steps of the patch selection method are illustrated in Figure 1. The initial step is to create a mask of the tissue on the slide. Most of the background can be removed by smoothing with a Gaussian filter and setting a threshold for the average intensity since the majority of the background is white. This threshold was set empirically to 90% of the maximum value. Further refinement is made in order to remove annotations on the slides that are typically blue or green. Finally, water stains and bubbles are removed by setting a lower bound on saturation, which was empirically set to 0.1.

**Fig. 1.**
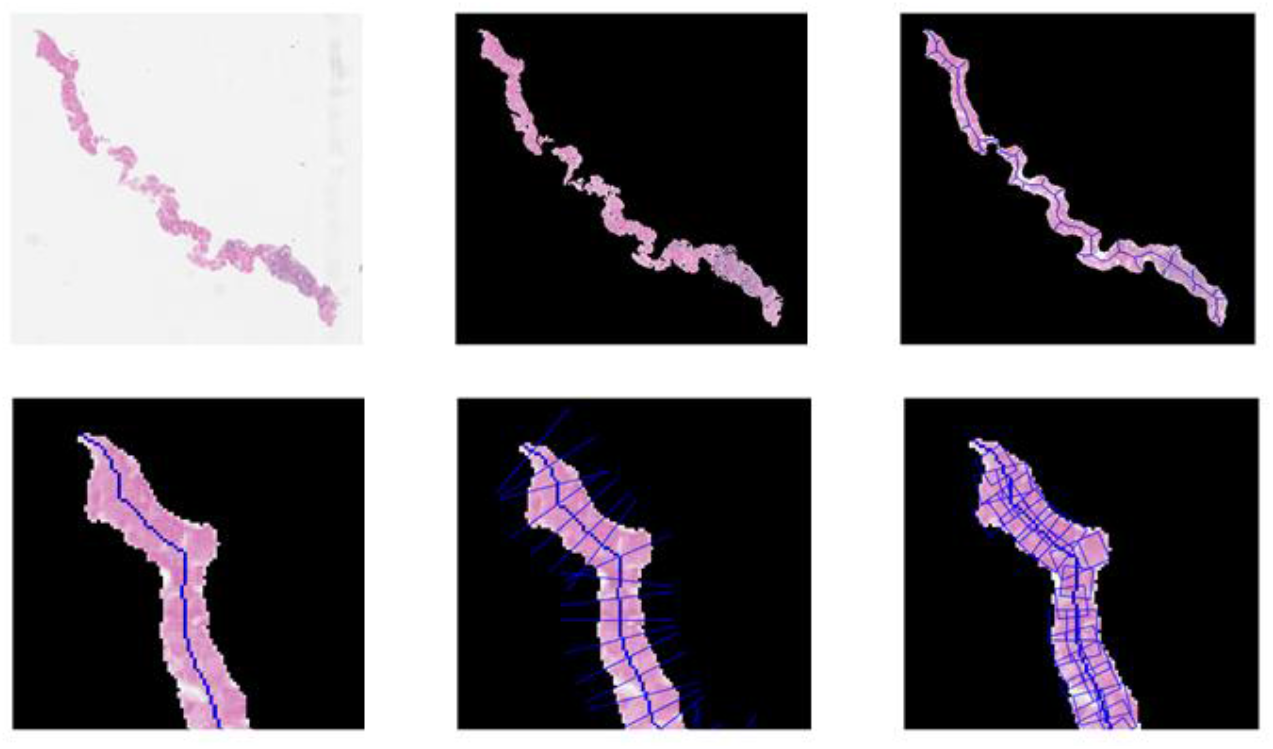
Outline of the proposed algorithm. The original slide (a) is smoothed and thresholded based on intensity and color (b). Morphological smoothing is performed and the image is skeletonized (c). Branches are pruned based on maximal geodesic distance (d) and perpendicular lines are found to the tangent at regular intervals (e). Patches are then selected along the perpendicular between the mask boundaries.

Once a tissue mask is found, it is smoothed using morphological closing. The skeleton of the smoothed mask is then found [11] and branches are removed by finding the endpoints with the maximum geodesic distance. The midline is then partitioned based on the patch size and overlap. In this study, a patch size of *x* × *y* = 512 × 512 pixels was used with an overlap of *s* = 25% as previously used in automatic Gleason grading [9]. Tangent lines are found at each of these locations by looking at the neighborhood of nine pixels along the midline and the perpendicular line is drawn until intersection with the mask boundary. A set of *n* patches are then chosen, equally spaced along the perpendicular line *l*, where

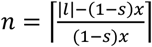

Finally, patches that intersect with less than 60% intersection with the mask are removed from the final set.

### 2.2 Path-RCNN Architecture

We leverage a previously trained framework for the cancer detection and Gleason score prediction in this paper. This network is a region-based convolutional neural network (R-CNN) framework for multitask prediction using an Epithelial Network Head and a Grading Network Head. It uses ResNet [12] as its backbone image parser. First, the image parser generates feature maps. These feature maps are then fed into two branches. In the left branch, it applies a two-stage procedure: the feature maps are first used by a Region Proposal Network (RPN) that generates region proposals (ROIs); second, a Grading Network Head (GNH) is used for predicting the class, box offset, and a binary mask for each ROI.

**Fig. 2.**
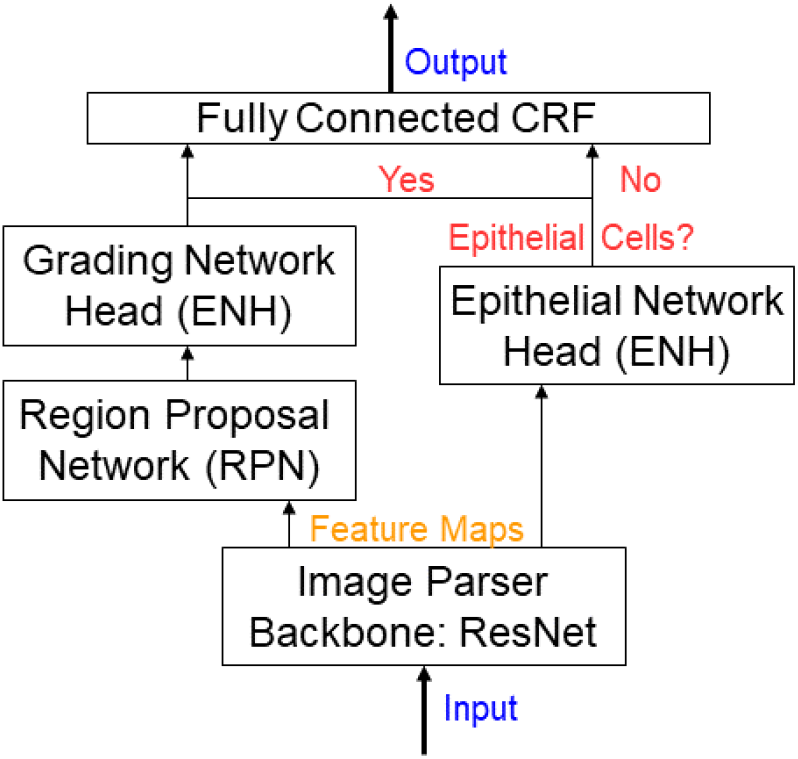
Overview of the Path R-CNN model architecture. ResNet was used as the backbone to extract feature maps from input images, which were fed into two branches in parallel. The first branch proposes regions for the grading network head to focus upon, which are then used to grade epithelial cell areas. The other branch determines if there is epithelial tissues in the image. In the final output, areas without epithelial cells are classified as stroma while other areas use the results from the grading network head.

In the right branch, the network outputs an epithelial cell score that detects the presence of epithelial cells in the image. We refer to this part as the Epithelial Network Head (ENH). The final prediction of the network depends on the results of the ENH and GNH. Finally, a post-processing step based on a conditional random field is applied to the prediction. Compared to a single task model, Path-RCNN can provide complementary contextual information, which contributes to better performance. Path-RCNN achieves state-of-the-art performance in epithelial cells detection and Gleason grading tasks simultaneously.

Models were implemented in Torch 7 [13] with two NVIDIA Titan Xp GPUs. Multiple separate data loading threads were used to accelerate training and testing. The average time required to generate a prediction mask for one tile was around 9 s.

### 2.3 Dataset and image preprocessing

The test dataset contains 59 prostate biopsy slides, each from a different patient. The scanning objective for all slides was set to either 20x (0.5 μm per pixel) or 40x (0.25 μm per pixel). Slides were manually segmented by a pathologist into four categories: benign/stroma, G3, G4, and G5. Because of the limited number of examples of G5 in our dataset, G4 and G5 were combined into a single class. For comparison, 512 × 512 tiles were extracted in the grid with 25% overlap. Tiles that contained less than 60% overlap with the tissue mask were removed. Both sets of tiles were normalized to account for stain variability [14].

Jaccard index, also known as the intersection-over-union (IOU), and overall precision (OP) were the method used for evaluation semantic segmentation in this study. IOU measures the extent of overlap between classification regions, while OP indicates the overall percentage of correct classifications. Significance is tested using Wilcoxon rank-sum tests with an alpha level of 0.05.

## 3 Results

The image-based patch selection method created slightly fewer patches per slide on average (140.9 vs. 142.1 for image-based and grid based selection, respectively), but the difference was not statistically significant (p=0.34). Nevertheless, the image-based method had significantly higher overlap with the image mask (p<0.001, see Fig. 3 left). Of the 59 biopsies in the dataset, 57 (96.6%) had higher IOU using the proposed image-based method, resulting in an average IOU of 0.78 compared to 0.72 for the grid-based method.

**Fig. 3.**
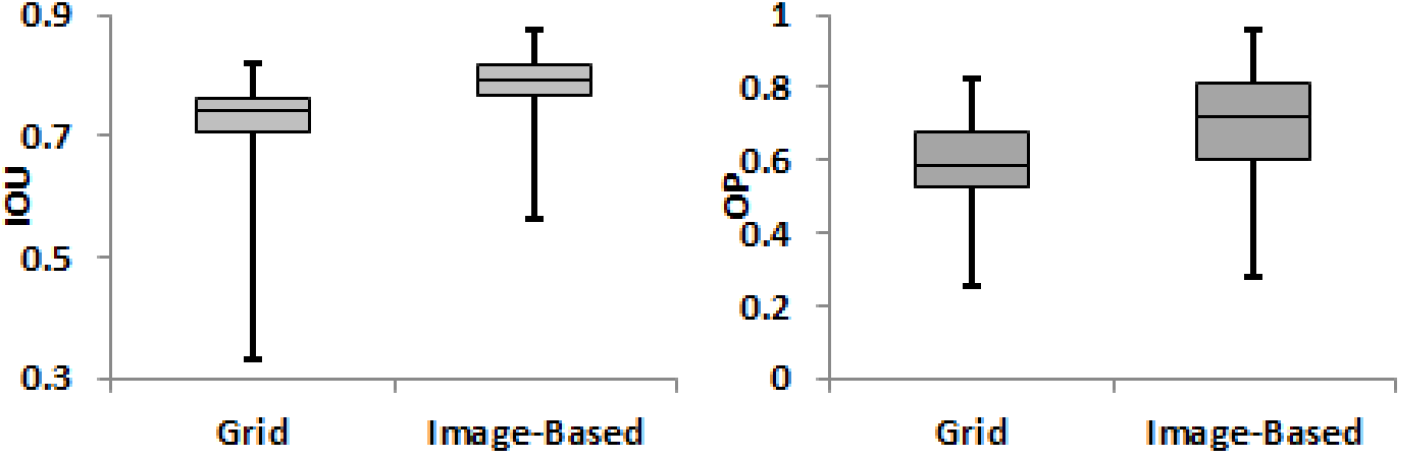
Intersection over union (IOU) of the area covered by the two patch selection methods comparted to the image mask (left) and overall classification precision (OP) across all pixels in a slide (right). The image-based method produced significantly better results in both cases (p<0.001).

The right side of Figure 3 shows the overall precision using the two patch selection methods. The classification precision improved for all of the slides in the test dataset when using the image-based patch selection method. The average IOU for each of the classes also improved on average when using the proposed method. This improvement was statistically significant for benign/stroma tissue and G3 cancer (p<0.001 and p=0.0028, respectively). It was not statistically significant, however, for high grade cancer (p=0.35).

Figure 4 displays the patches selected for several examples. In cases where tissue boundaries do not align with patch boundaries, the grid-based method includes significant white space in patches selected. There are also areas where tissue is not covered by the tile grid because alignment caused tiles to fail to meet the tissue threshold for inclusion in the analysis. The proposed method had some missing coverage, particularly around sharp curves and areas where the tissue did not have defined directionality.

**Fig. 4.**
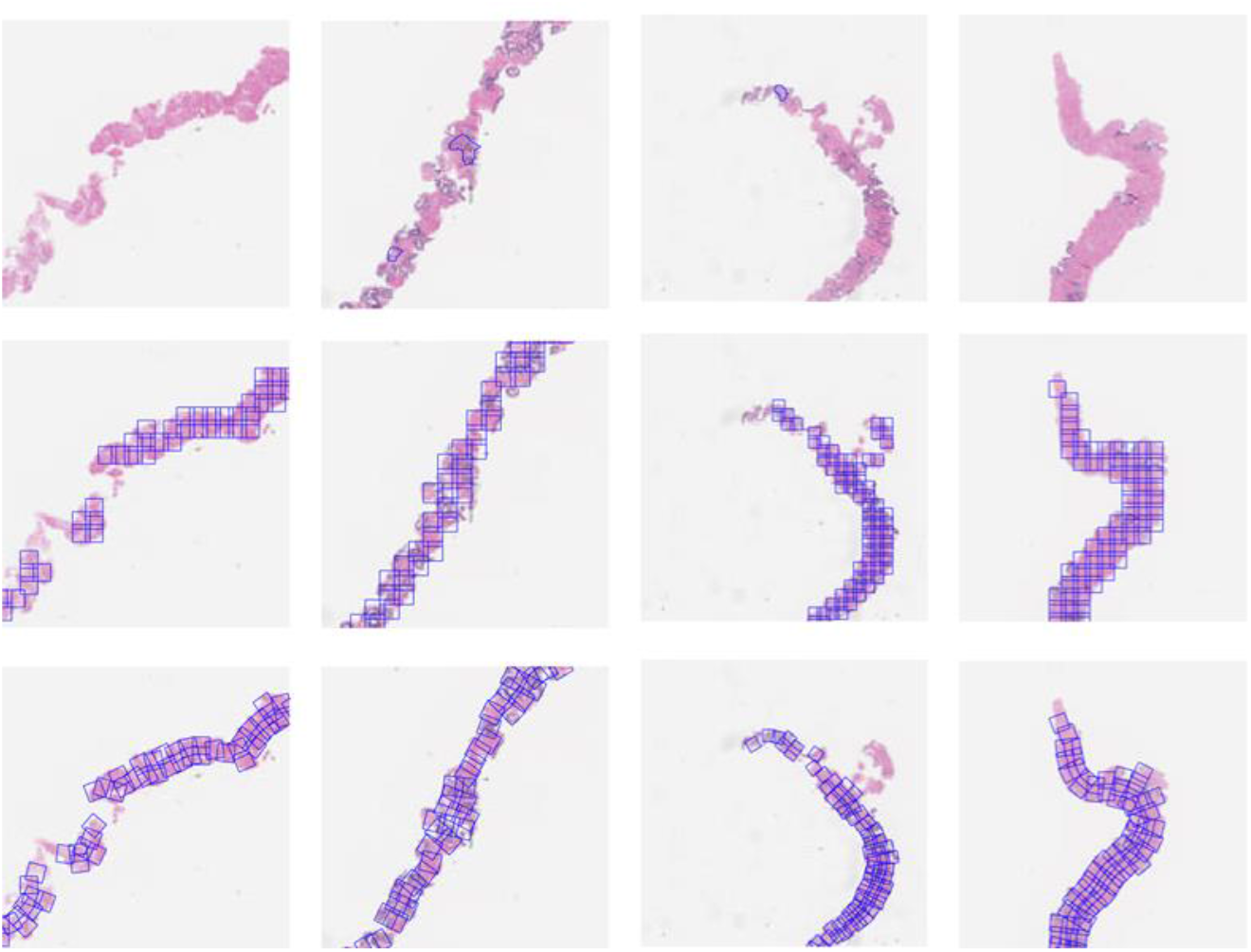
Examples of patch selection using the two algorithms. Original images are shown on the top row. The second row shows patches extracted using a grid-based method. The bottom row shows the results using the proposed method.

## 4 Discussion

The proposed image-based patch selection method produced fewer patches on average than the grid based approach, but the patches provided better coverage of the biopsy mask. The improved coverage results from two factors. First, the rotation of the patches allows them to line up with the boundary of the prostate biopsy, reducing the amount of white space that is included (see Fig. 4 column 2). Second, by removing the restriction that patches fall within a grid, thin segments that fall on the grid boundary can be sampled, which they might be removed from grid-based approaches because of insufficient tissue in each grid location (see Fig. 4 column 1).

Using the image-based patch selection method, classification performance improved significantly in terms of both overall precision and IOU. This is improvement is largely due to the reduction in patches that lie on the tissue boundary and contain significant amounts of white space. In addition, patches found using the proposed method were less likely to miss regions of the biopsy, which could potentially contain cancer tissue. The difference in IOU value for G4/G5 detection was not statistically significant, however. This is largely because of the lack of high grade cancer in the dataset. Only 34% of the subjects tested had G4 or G5 annotations. A more balanced testing set might yield significant improvements in G4/G5 detection as well.

The proposed method demonstrates significant performance above traditional grid selection methods. However, there are some limitations that could be improved in future work. The largest limitation is that patch selection oversamples on the inside of curves and undersamples on the outside (see Fig. 4 column 3). This phenomenon occurs because sampling frequency along the length of the biopsy is determined based on the midline of the mask. A possible improvement that would address this problem would be to evaluate the sampling frequency separately for patches that are beyond a set distance from the midline. Another limitation is that some parts of a biopsy can be missed if the biopsy bifurcates or folds so that the directionality is unclear (see Fig. 4 column 4). Allowing for multiple branches in the biopsy could alleviate this problem.

While the analysis presented here was restricted to providing patches for deep learning applications, the methods described here can be used for other clinical applications as well. One possible application would be for integrating histopathological information with radiographic imaging. Multiparametric MRI can provide the locations of biopsy needles within the prostate, which allows for comparison and integration of pathological findings with radiological evaluations [15]. However, annotations and segmentations within the prostate slide cannot be localized within the MR image because there is not a direct mapping from pixels within a slide to the MR volume. Using the method proposed here, annotations can be aggregated along the length of the biopsy, which can then allow these values to be associated with locations along the biopsy needle in MR space.

## 5 Conclusion

The proposed image-based patch selection algorithm increases the coverage of deep learning classifiers for histopathological slides. Because patches contain more context, pixel classification accuracy significantly improves when using the proposed method. Implementing patch selection strategies could significantly improve the potential for a deep learning based automatic cancer detection system.

## Acknowledgements

The authors would like to acknowledge support from the UCLA Radiology Department Exploratory Research Grant Program (16-0003), NIH/NCI R21 CA220352, and NIH/NCI 5P50CA092131-15:R1. KVS acknowledges support from an AMA Foundation Seed Grant, NIH NCI F30CA210329, NIH NIGMS GM08042, and the UCLA-Caltech Medical Scientist Training Program. We gratefully acknowledge the support of NVIDIA Corporation with the donation of the Titan Xp GPUs used for this research.

